# Diversity and evolution of the human anellome

**DOI:** 10.1101/2024.05.13.593858

**Authors:** Sejal Modha, Joseph Hughes, Richard J. Orton, Spyros Lytras

## Abstract

Anelloviruses are a group of small, circular, single-stranded DNA viruses that are found ubiquitously across mammalian hosts. Here, we explored a large number of publicly available human microbiome datasets and retrieved a total of 829 anellovirus genomes, substantially expanding the known diversity of these viruses. The majority of new genomes fall within the three major human anellovirus genera: *Alphatorquevirus*, *Betatorquevirus* and *Gammatorquevirus*, while we also present one new genome of the under-sampled *Hetorquevirus* genus. The phylogeny reconstructed from the conserved ORF1 gene reveals three additional, previously undescribed, human anellovirus clades. We performed recombination analysis and show evidence of extensive recombination across all human anelloviruses. Interestingly, more than 95% of the detected events are between members of the same clade and only 15 inter-clade recombination events were detected. The breakpoints of recombination cluster in hotspots at the ends and outside of the ORF1 gene, while a recombination coldspot was detected within the gene. Our analysis suggests that anellovirus evolution is governed by homologous recombination, however events between distant viruses or ones producing chimaeric ORF1s likely lead to non-viable recombinants. The large number of genomes further allowed us to examine how essential genomic features vary across anelloviruses. These include functional domains in the ORF1 protein and the nucleotide motif of the replication loop region, required for the viruses’ rolling-circle replication. A subset of the genomes assembled in both this and previous studies are completely lacking these essential elements, opening up the possibility that anellovirus intracellular populations contain defective virus genomes (DVGs). Overall, our study highlights key features of anellovirus genomics and evolution, a largely understudied group of viruses whose potential in virus-based therapeutics is recently being explored.

## 1 Introduction

Advances in high throughput sequencing technologies and reduced sequencing costs have had a strong impact on virus research specifically in the context of virus discovery (Wolf et al., 2020). Viruses are the most abundant entities in our ecosystem, and they are found in all environments captured and catalogued by researchers (Edwards and Rohwer, 2005; Nayfach et al., 2021). Metaviromic approaches are less biased compared to traditional culture-based techniques and have enabled the discovery of a large, and still expanding, diversity of previously unknown viruses (Edgar et al., 2022).

Anelloviruses are small, circular, single-stranded DNA (ssDNA) viruses with negative-sense genomes that range from 1.6-3.9 kb in length (Miyata et al., 1999; Varsani et al., 2021). The first identified anellovirus, named torque teno virus (TTV), was found in 1997 in the blood sample of a hepatitis patient (Nishizawa et al., 1997). Since then, investigators hypothesised the potential involvement of these viruses in disease, partly because of their prevalence in immunocompromised patients and blood transfusion samples (Itoh et al., 2000; Simmonds et al., 1998). However, it now seems that anelloviruses are not pathogenic and are commensals instead, ubiquitously found in virtually all individuals (Cossart, 2000; Vasilyev et al., 2009; Koonin et al., 2021). For years, the non-pathogenic nature of these viruses has made them fall outside of the focus of mainstream virology research and many aspects of their genomics, diversity and overall biology remain poorly understood. The recent push for ramping up virus discovery through metagenomics and the potential use of anelloviruses in viral-based therapeutics has re-ignited the interest in anellovirus research (Arze et al., 2021; Liou et al., 2022; Butkovic et al., 2023).

In the last few years, a large diversity of anelloviruses has been catalogued in a range of studies led by Moustafa et al. (2017), Cebriá-Mendoza et al. (2021), Tisza et al. (2020) and Arze et al. (2021). These genomes have been found in closely monitored blood transfusion patients, but also in metagenomic datasets from many other types of samples including faeces, nasal secretions, saliva, urine, and bile, suggesting their omnipresence in all types of human microbiomes and non-specific tissue tropism (Kaczorowska and Hoek, 2020). Their presence is not unique to humans, with diverse anelloviruses having been discovered in many other mammals (Okamoto et al., 2000; Abe et al., 2000; H. Okamoto et al., 2001; Fahsbender et al., 2017; Kraberger, Opriessnig, et al., 2021). There is currently a total of 156 species classified in 30 genera under the *Anelloviridae* family according to the International Committee on Taxonomy of Viruses (ICTV) Master Species List v38 (ratified in September 2023). The higher taxonomic placement of the family has been ambiguous until recently due to the viruses’ genetic distinctness to other circular DNA virus groups. However, Butkovic et al. (2023) propose that anelloviruses are classified into a new phylum, the ‘Commensaviricota’, under the *Shotokuvirae* kingdom (realm *Monodnaviria*). The human-infecting members of the family are classified into four genera: *Alphatorque-virus* (TTV), *Betatorquevirus* (TT mini viruses - TTMV), *Gammatorquevirus* (TT midi viruses - TTMDV), and *Hetorquevirus* (Varsani et al., 2021). The three major genera of human anelloviruses have different genome lengths. TTVs have the largest genomes that range from 3.6 to 3.9 kb, TTMVs have genomes that range between 2.8 to 2.9 kb whereas TTMDVs have a slightly wider genome size range from 2-3.2 kb (Taxonomy of Viruses, 2011). All anellovirus genomes code for one major open reading frames (ORFs), referred to as ORF1. Some genomes also contain few partially overlapping accessory ORFs, but their function is less well characterised and will not be analysed in this paper (Kaczorowska and Hoek, 2020). ORF1 is used for species and genus delineation and has recently been confirmed to function as the capsid protein of the virion, containing a jelly-roll-like domain and forming a capsid of 60 ORF1 monomers in an icosahedral (T=1) conformation (Liou et al., 2022; Butkovic et al., 2023). Another distinct feature of the ORF1 is an arginine (and lysine)-rich, N-terminal sequence motif (ARM) that localises within the capsid polymer. This positively charged part of the protein likely stabilises the genome when inside the virion and may be involved in genome replication and packaging (Liou et al., 2022). Like most circular DNA viruses, anelloviruses replicate their genome through rolling-circle replication. The non-coding part of the genome contains a small (eight nucleotide), conserved sequence motif, surrounded by GC-rich flanking regions, which is likely cleaved to act as the origin of replication (Villiers et al., 2011). Lastly, much of the sequence diversity observed across the anelloviruses may be caused by extensive recombination, taking place during the viruses’ replication (Worobey, 2000; Arze et al., 2021; Kraberger, Serieys, et al., 2021).

In a previous study by Modha et al. (2022) Unxplore - a systematic framework to identify, assemble, and quantify unknown sequences present in various human microbiomes - was developed and applied to a range of human microbiome datasets. Modha et al. (2022) examined 40 different studies covering 963 samples across 10 human microbiomes, such as faecal, oral, and skin, and found that 2% of sequences remain unclassified, indicating a vast potential for discovering new viruses. In this study, we extend these findings to mine sequences categorised as known and partially known in Modha et al. (2022). Specifically, we analyse, identify, and focus on the diversity of anelloviruses present in the publicly available human microbiome datasets (n=2,596) that were investigated using Unxplore (Modha et al., 2022), expanding our understanding of the anelloviruses circulating in humans - collectively referred to as the human ’anellome’. We shed light on the phylogenetic relatedness between the human anellovirus genera, their overall patterns of genomic recombination, and the distribution of essential features in their genomes.

## 2 Results and Discussion

### 2.1 A wide diversity of *Anelloviridae* in human metagenomes

We applied Unxplore - a previously developed framework for assembling uncharacterised contigs - on 2,596 publicly available human microbiome datasets and implemented an array of virus contig detection methods to look for anellovirus-related sequences (Modha, 2022). In this way, we retrieved a total of 829 contigs representing distinct anellovirus genomes and encoding complete ORF1 genes. We combined this dataset with all published human anellovirus ORF1 coding sequences and used the resulting coding sequence alignment to infer a maximum likelihood phylogenetic tree encompassing the broadest human anellovirus genome set to date (n=2,605; Figure 1A). The tree formed three major clades, consistent with the three largest anellovirus genera: *Alphatorquevirus*, *Betatorquevirus* and *Gammatorquevirus*. The largest number of ORF1 sequences assembled in this study (418) are members of the *Alphatorquevirus* genus which was also the taxon with the most known isolates before this study. Interestingly, we identified 228 members of the *Gammatorquevirus*, one of the previously least sampled genera, followed by 170 *Betatorquevirus* sequences and 1 *Hetorquevirus* genome. The latter is the most recent anellovirus genus found in humans with only two previously known members. Twelve sequences assembled in this study do not fall within the 4 previously defined taxonomic clades, forming two distinct clusters in the phylogeny we refer to as clades X1 and X2 respectively (Figure 1A). Based on the current sampling, the ORF1 gene of a genome from a different study (NCBI accession: MH649127) also fell outside of the main clades, forming a lone branch that we refer to as clade X3. These three clades could represent novel genera within the *Anelloviridae* and should be considered in the future taxonomic assessments of the family.

**Figure 1:**
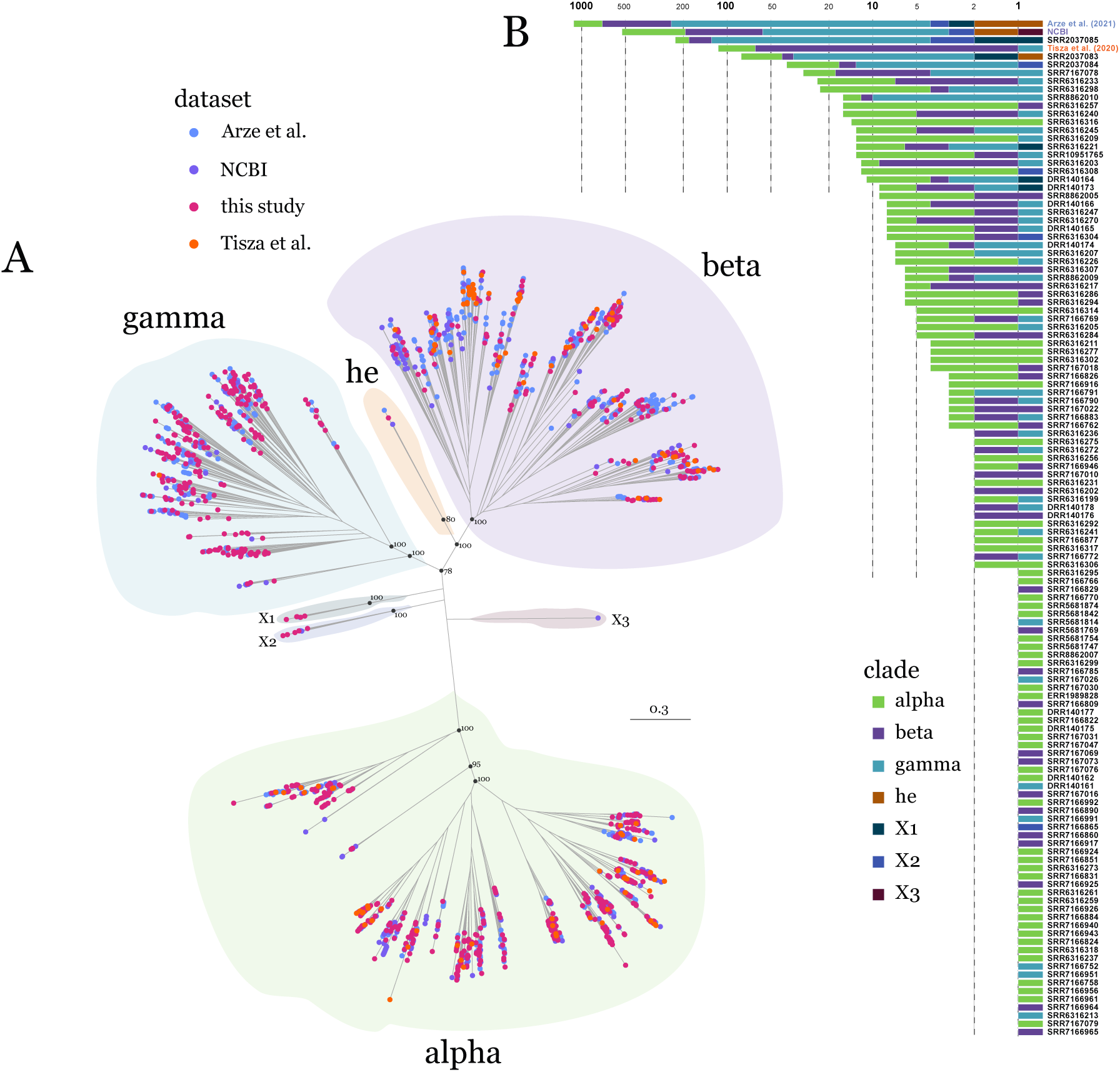
Phylogenetic and dataset distribution of all known human anelloviruses. A) Unrooted phylogenetic tree of all available anellovirus ORF1 coding sequences. Tips are coloured by the dataset the sequences were derived from. Shading indicates the taxonomic classification of distinct clades in the tree. Ultrafast bootstrap support values are shown for key basal nodes. B) Stacked bar plot showing the number of ORF1 sequences retrieved from each SRA sample (in this study) and from other existing datasets, separated by the sequences’ clade definition (consistent with panel A). Bars are ordered by the total number of sequences in each sample/dataset. The x axis is in a base 10 logarithmic scale, absolute numbers corresponding to each axis line are shown as axis labels.

The larger proportion of *Gammatorqueviruses* (27.5% of the genomes) was the clearest discrepancy between our dataset and that of previous studies on human anelloviruses (Figure 1B). Surprisingly, the vast majority of genomes, including more than half of our dataset’s *Gamma-torqueviruses*, were found in a single SRA sample (SRR2037085, bioproject: PRJNA230363) (alpha: 44, beta: 55, gamma: 124, X1: 2, X2: 2) associated with human oral microbiome samples (Wang, Gao, et al., 2016; Wang, Jia, et al., 2020). We further investigated the samples from this study to identify potential factors explaining this pattern. Unlike the majority of other metagenomic samples deposited by the same study, which were derived from individual patients, sample SRR2037085 contained pooled salivary samples of five individuals that were subsequently amplified for DNA viruses by purification of virus-like particles (VLP). We expect that the downstream processing steps unique to this sample are part of the reason why so many anelloviruses were retrieved from its sequencing data. VLP purification is likely a valuable step when preparing sequencing samples for the identification of anelloviruses and other small DNA viruses. Consistently, the next two samples with the largest number of assembled genomes in our dataset have undergone the same processing steps (SRR2037083 and SRR2037084; Figure 1B). This finding may also imply that many diverse anelloviruses are missed from the metagenomic datasets when using conventional DNA sequencing protocols.

Even though this amount of, what seems to be, intra-host genetic diversity may be surprising for many viruses (especially those causing acute infection), similar findings have been shown before for anelloviruses. Specifically, Arze et al. (2021) quantified the number of distinct anellovirus lineages in individual blood donors and recipients and found up to approximately 90 and 300 lineages in a single donor and recipient respectively. Contamination unique to the SRR2037085 sample cannot be excluded, but all genomes assembled from it are distinct and dispersed across the phylogeny of other human anelloviruses, making a single contamination source highly unlikely. Finally, to assess whether the large number of genomes was a technical artefact of our sequence assembly methodology, we reassembled the sample’s reads using a second assembly algorithm and confirmed the presence of most distinct genomes in the sample (see Methods).

Since the microbiome datasets come with information about the type of tissue the samples came from, we questioned whether there were specific patterns of microbiome clustering in the ORF1 phylogeny. This does not seem to be the case with no apparent clustering in the microbiome source across any of the clades in the tree (Figure S1). The majority of viruses were retrieved from blood samples (2028/2605), however this likely reflects the sampling bias towards blood transfusion patients (Arze et al., 2021). If anelloviruses circulate ubiquitously in the blood it is also difficult to exclude tissue contamination from circulating blood or injuries in the other sample sources assessed here (oral, pulmonary and serum microbiomes). Based on the current metagenomic data available there is no evidence to suggest that different human tissues harbour a distinct diversity of anelloviruses.

### 2.2 Extensive recombination happens non-randomly across the genome

The wide diversity we observe across the human anelloviruses could be partly due to the viruses frequently recombining with one another. We used an array of recombination detection methods on the full genomes available in our datasets (n=1,472) to explore the extent of this process present in the evolution of human anelloviruses. We detected 345 unique recombination events, confidently inferred by at least 3 independent detection methods. Strikingly, 95.7% of these recombination events (330) were between genomes of the same genus (intra-lineage) with only 15 (4.3%) events having probable parental sequences from different genera (inter-lineage). It is worth noting that intra-lineage recombination events are expected to be more difficult to detect because the parental sequences are more genetically similar to one another. Hence, we find that recombination happens extensively between closely related human anelloviruses, however interlineage recombinants are rarely observed in the available genomes. Many sequencing samples contained a mixture of anelloviruses from different clades (Figure 1B), suggesting that the lack of inter-lineage recombination events is not due to different clades infecting different tissues or individuals. Instead, the most likely explanation would be that inter-lineage recombinants are non-replicative due to genomic incompatibilities. Among the intra-lineage recombination events detected, there are 241 within-alpha, 48 within-beta and 41 within-gamma. These proportions are consistent with the number of genomes from each clade included in the recombination analysis (alpha: 803, beta: 370, gamma: 283). This implies that including more genomes in the analysis is likely to increase the number of detectable intra-lineage recombination events.

We further investigated whether breakpoints of recombination were clustering in specific parts of the genome. We used the breakpoint distribution test (BDT) and the recombination range test (RRT) with the breakpoints from all 345 recombination events detected by our original analysis. These tests compare the number of breakpoints detected in each window across the genome alignment to the informative nucleotide sites in that window, determining the likelihood of detecting a breakpoint in this region of the alignment. Both tests show a very similar pattern, where most of the non-coding part of the genome has more breakpoints than expected by chance, being recombination hotspots (Figure 2). Interestingly, the replication loop region is also determined as a hotspot of recombination. In contrast, the majority of the ORF1 coding region is a clear coldspot of recombination, with very few detected breakpoints within it (Figure 2). It is worth noting that the 5’ and 3’ ends of the coding region do contain distinct recombination hotspots. The lack of observable recombination breakpoints within ORF1, despite the extensive amount of recombination detected across the rest of the genome, may indicate that functional constraints within the gene make chimaeric ORF1s mostly deleterious for further virus replication. Another explanation might be related to breakpoints being more likely to occur closer to the origin of replication (the replication loop). These two hypotheses are not exclusive of one another. The overall pattern of recombination hot- and coldspots across the human anellovirus genomes is strikingly consistent with the results of similar analysis on felid anelloviruses (Kraberger, Serieys, et al., 2021), suggesting that the mechanisms behind anellovirus recombination are conserved across the anelloviruses and not determined by the host they infect. Paired with the lack of inter-lineage recombination events, we show how distinct restrictions apply to recombination across the *Anelloviridae*, both in terms of which viruses can recombine and where in the genome the recombination breakpoints can take place.

**Figure 2:**
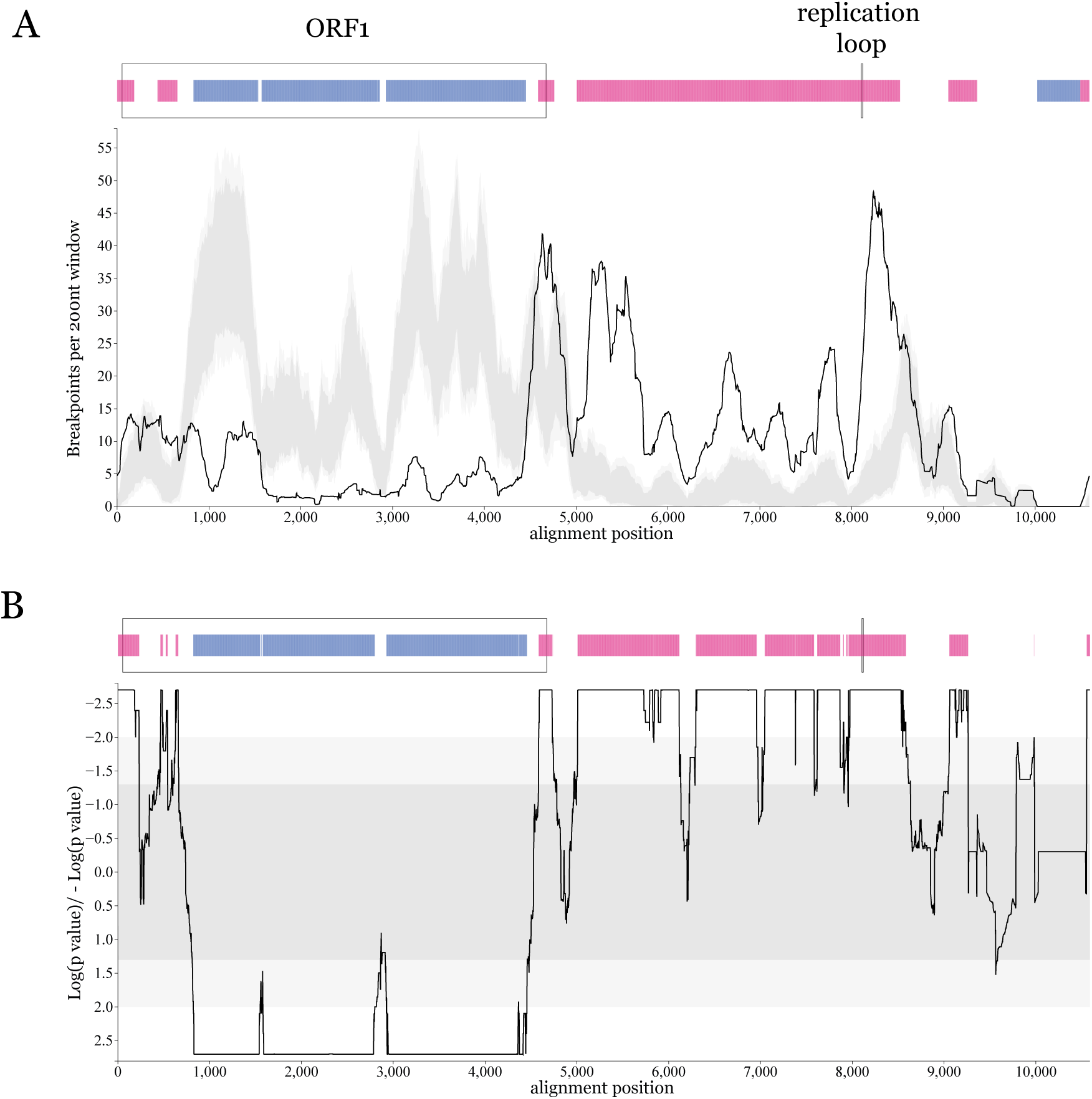
Hotspots and coldspots of recombination across human anelloviruses. Distribution of recombination hot- and coldspots across the alignment based on the RRT (A) and the BDT (B) methods. For both plots, dark grey shading represent 95% and light grey shading 99% confidence intervals of expected recombination breakpoint clustering under random recombination. Peaks above the shaded area represent recombination hotspots and drops below represent coldspots, annotated on the corresponding genome schematic above each plot by vertical pink and blue lines, respectively (99% confidence). The regions of the alignment corresponding to the TTV ORF1 coding sequence and replication loop are annotated on both genome schematics, labelled on panel A.

### 2.3 Genus-specific genomic features

The novel genomes we provide in this study also allows us to explore the diversity of key genomic features in the human anelloviruses. The N-terminal region of the anellovirus ORF1 protein is characterised by a very high content of positively charged residues (primarily arginines and lysines; Figure 3D). This distinct protein feature is referred to as the arginine-rich motif (ARM) and has been shown to be involved in genome packaging by forming binding interactions with the viral DNA (Liou et al., 2022). We quantified the positively charged content of this motif for each ORF1 protein in our dataset by counting the number of arginine (R) and lysine (K) residues in the N-terminal part of the protein, corresponding to the ARM region. Based on the distribution of R/K residues present across all the ORF1 ARM regions analysed here, we proceeded to accept regions with 20 to 60 R/K residues as complete ARMs, used for the downstream comparisons (see Methods, Figure S2). We find a clear separation in the number of R/K residues by the virus genus, consistent with the genome size of each taxon. *Alphatorquevirus* members had the largest number of R/Ks (mean = 48), followed by *Gammatorquevirus* (mean = 40), with the shortest taxon (*Betatorquevirus*) having the least R/K residues (mean = 33) (Figure 3A). 248 ORF1 sequences (9.5% of the dataset) had less than 10 R/K residues in their N-terminal region, most having no R/Ks at all, and were deemed to have a partial or missing ARM region (Figure S2).

**Figure 3:**
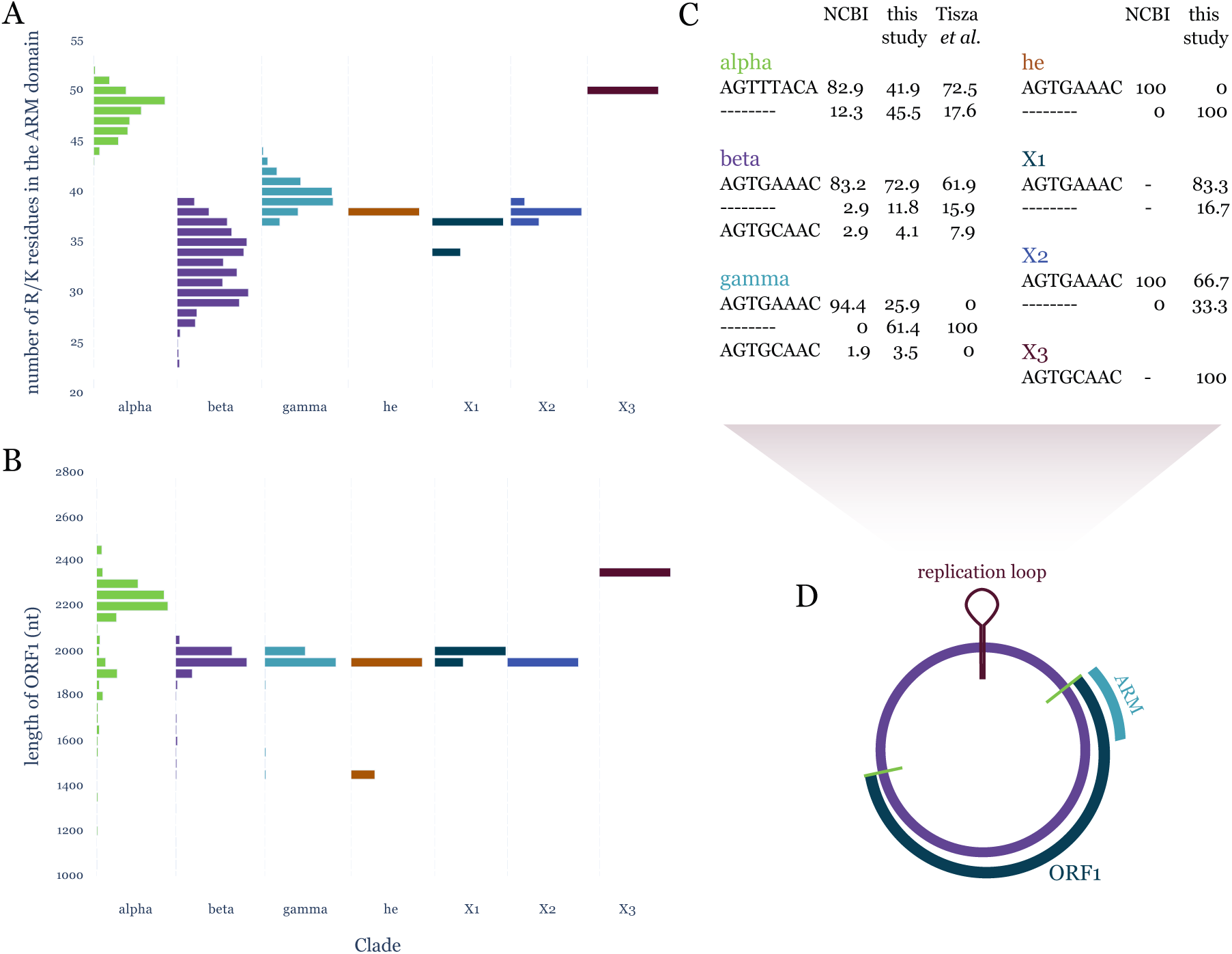
Diversity of key genomic features between human anellovirus groups. A) Distribution of the content of arginine (R) and lysine (K) residues in the ARM domain defined for each ORF1 sequence, separated by phylogenetic group. Only sequences with a number of R/K residues between 20 and 60 are used for this analysis (see Methods, Figure S2). Bar heights are normalised by the number of genomes in the largest bar. B) Distribution of ORF1 coding sequence lengths for all ORF1s used in this study, separated by phylogenetic group. Bars represent bins of 50 nucleotides of ORF1 length each. Bar heights are normalised by the number of genomes in the largest bin. C) Proportion of major replication loop genotypes (present in >2% of the sequences in each group) for genomes of each phylogenetic group, separated by the dataset the sequences were retrieved from. Only ORF1 coding sequences were available for the Arze *et al*. (2021) dataset, so it does not appear in this analysis. D) Schematic of the anellovirus genome, highlighting the relative location of the key genomic features.

Another key feature of the anellovirus genome is the conserved replication loop believed to be cleaved and subsequently act as the origin of rolling-circle viral replication (Figure 3D). We confirm that replication loop sequences are clearly conserved within anellovirus groups (Figure 3C). As previously described, beta, gamma, and he anelloviruses have the same, conserved AGTGAAAC replication loop motif, however about 2-8% of the beta and gamma genomes across datasets also contain the less prevalent AGTGCAAC motif. This single nucleotide difference variation in this region is further reflected in the sequences of the three ‘X’ clades described here with X1 and X2 viruses having AGTGAAAC and the X3 clade genome having the less prevalent AGTGCAAC motif. The alpha clade has a distinct replication loop to all other human anellovirus clades, AGTTTACA, which is consistently conserved within the clade (Figure 3C). Along with the longer branch separating the alpha clade from the other groups in the ORF1 phylogeny (Figure 1A), *Alphatorqueviruses* have likely separated from the other groups earlier in the viruses’ evolutionary history.

In addition to the conservation of replication loop sequences, we observed that many of the genomes assembled both in this study and by previous research completely lack the replication loop (Figure 3C). The proportion of replication loop presence varied notably across datasets, from 96.3% of the gamma genomes retrieved from NCBI containing a complete replication loop to only 41.9% of alpha genomes and 29.4% of gamma genomes reported in this study having the loop sequence. The variation in the presence of a replication loop in the assembled genomes between clades and studies suggests that it cannot be fully explained by the DNA sequencing quality of each dataset or the assembling methodology in each study.

The lack of the replication loop region and the ARM-encoding part of the ORF1 gene in many of the available anellovirus genomes would imply that these genome sequences are missing key features for the viruses’ replication. If this incompleteness were an artefact of sequencing read quality then we would expect the partial contigs to have lower overall read coverage compared to the complete ones. This is not the case, at least in our dataset, since the contig coverage distribution is identical between contigs with and without replication loops and ARM regions (Figure S3). Another indication of the missing features being a sequencing or assembly artefact would be a read coverage dropout near these genomic regions. We inspected the per site coverage of example contigs with high mean coverage and absence of the ARM region and find that coverage is generally uniform across contigs with no evidence of local dropouts (Figure S4).

These observations open up the possibility of these seemingly partial genomes being intact DNA molecules present in the samples. Villiers et al. (2011) were the first to experimentally validate the presence of diverse ’subviral’ anellovirus genomes of varying lengths when the viruses were cultured *in vitro*. A recent study by Kaczorowska, Timmerman, et al. (2023) monitored long-term evolution of distinct anellovirus lineages in infected individuals and further confirmed the existence of dynamic heterogeneous intra-host genomic populations. Consistent with these studies, we propose the possibility that anellovirus-infected cells can contain diverse genetic populations composed of intact, replicative genomes as well as defective viral genomes (DVGs), some of which may be non-replicative. Other than complete genomes, metagenomically assembled contigs will likely also reflect DVGs that comprise the overall genomic heterogeneity in each sample.

## 3 Conclusions

In this study, we provide 829 novel anellovirus genomes retrieved from more than 2,000 publicly available human metagenomic samples. These viruses are widely distributed across the known diversity of human anelloviruses, reflecting the heterogeneity of genomes within the analysed metagenomes. We use all known human anellovirus genomes to study the patterns of recombination in this virus group. Our analysis reveals that, even though recombination takes place extensively between genomes, there are restrictions pertaining to the mechanism of anellovirus recombination. Specifically, the vast majority of detected recombination events are between viruses belonging to the same phylogenetic clade, while breakpoints of recombination rarely occur within the ORF1 coding region. The absence of otherwise highly conserved genomic features necessary for virus replication in some of the assembled genomes leads us to hypothesise that anellovirus populations potentially include substantial proportions of DVGs. We expect that this intra-host heterogeneity is a major driver of the extensive recombination patterns we describe in this manuscript. The genomic diversity we present here is likely only the tip of the iceberg that is the human anellome. Anelloviruses have recently shown promise to be used as biomarkers of immunosupression in patients (Castain et al., 2024) as well as vectors for gene therapy (Prince et al., 2024). Better describing the diversity of these ubiquitous commensals can improve the development of medical applications, but also reveal more on how circular DNA viruses utilise features like recombination and DVGs during their evolution.

## 4 Methods

### 4.1 Metagenomic assembly and virus sequence identification

61 studies spanning 11 human microbiomes (bodily sites/sample types) were analysed as part of a bigger project looking for signatures of unknown sequences in human microbiome samples using the UnXplore framework developed by (Modha et al., 2022). Briefly, the UnXplore framework is an unknown sequence identification and quantification framework that can assemble high-quality contigs from human microbiome datasets; although focussed on ’unknown’ sequences, the framework identifies those of known origin at the same time. We utilised the samples analysed as part of (Modha et al., 2022) (n=963) as well as an additional 2,596 human blood microbiome datasets retrieved from the Sequence Read Archive (SRA) repositories. Each sample was *de novo* assembled and investigated for the presence of unknown sequences (often referred to as biological “dark matter”) using the UnXplore framework.

The contig set generated from the UnXplore analysis was consolidated and filtered to remove short contigs that were <1kb long. A total of 7,196,090 contigs were then submitted to three separate virus prediction tools: VirSorter2, DeepVirFinder and TetraPredX (Guo et al., 2021; Ren et al., 2020) (https://github.com/sejmodha/TetraPredX). The prediction results were filtered using the following criteria. For DeepVirFinder, contigs with a score >=0.9 and p-value <0.05 were selected, for VirSorter2 all predicted contigs with a minimum score 0.5 were selected and for TetraPredX all contigs with viral signal probability >=0.95 and all other class probability <=0.5 were selected. These contigs were then searched against an extensive set of nucleotide (nt) and protein sequence (refseq_protein) databases for further validation. The top 25 hits for each contig were extracted and the Lowest Common Ancestor (LCA) was computed from the sequence similarity results obtained using BLAST (nucleotide) (Camacho et al., 2009) and DIAMOND (protein level) (Buchfink et al., 2021). Based on the LCA, each contig’s superkingdom was retrieved from the NCBI taxonomy database using ete3 (Huerta-Cepas et al., 2016). A proportion of virus hits and subsequently virus families were determined for each contig using the ‘ExtractLCA.py’ and ‘ExtractLCABLASTM6.py’ scripts included in UnXplore.

In total, 272,827 contigs of interest (with LCA root or viruses) were examined which includeda set of 122,884 confirmed virus contigs. From this set, a total of 2,005 contigs were confirmed and matched exclusively to anelloviruses and were extracted. This set included contigs that were 1,000-8,220 bases long. From this set, contigs that were shorter than 2 kb (n=709) and longer than 4 kb (n=19) were removed based on the known length of anellovirus genomes. The remaining 1,277 contigs that were between 2-4kb long were retained and used for all subsequent analyses described in this study.

To validate the assembly of contigs from sample SRR2037085 from which we retrieved an unusually large number of anellovirus genomes, we also assembled the reads in this sample with MEGAHIT (Li et al., 2015), a different assembler made specifically for metagenomic assembly. We extracted MEGAHIT contigs with lengths of more than 300 nucleotides and used BLAST (Camacho et al., 2009) to compare them to the SRR2037085 contigs assembled from our original pipeline. We confirm that 269 contigs assembled with MEGAHIT had matches to the reference contigs with at least 90% coverage.

All anellovirus contigs are available publicly and can be accessed using BioProject PR-JEB75196 [*upon publication*].

### 4.2 Processing of virus genomes

To obtain a set of representative sequences, 1,277 anellovirus contigs were clustered using MM-Seqs2 easy-cluster pipeline with –min-seq-id 0.95, –cov-mode 1 -c 0.99 parameters (Steinegger and Söding, 2017), and a set of 840 representative sequences was obtained. To ensure that an ORF1 protein is encoded by each of the sequences we used the EMBOSS getorf application (Rice et al., 2000) with a minimum size of 1000 nucleotides and assuming that the sequences are circular. We could not identify an ORF1 using these criteria for 11 sequences which were excluded in downstream phylogenetic analysis, resulting in a final set of 829 novel anellovirus genomes.

To complement the novel genomes presented in this study with the available anellovirus diversity, we retrieved all human anellovirus genome sequences discovered in a study led by Tisza et al. (2020) (BioProject: PRJNA396064), as well as those analysed in a recent study led by Varsani et al. (2021). Getorf was also used in these sequences, as above, to identify the coordinates of the ORF1 coding region. All genomes were then rotated so that the ORF1 start codon was at the start of the sequence to enable downstream sequence aligning. Genome sequences with no identifiable ORF1 coding sequence were excluded.

Another large-scale anellovirus discovery study led by Arze et al. (2021) was recently published, however, only ORF1 coding sequences were made available by the authors. We incorporated these sequences (downloaded from https://github.com/ring-therapeutics/anellome_ paper) into our dataset and downloaded the full genome sequences for any NCBI genomes analysed by Arze et al. (2021) that were not already included. This led to a final whole-genome dataset consisting of 1,472 sequences (this study: 829, NCBI: 528, Tisza: 115) and an ORF1 dataset of 2,605 sequences (including an additional 1,133 ORF1 sequences from Arze et al. (2021)).

### 4.3 Phylogenetic analysis

We aligned all 2,605 ORF1 protein sequences using mafft (–genafpair option) (Katoh and Standley, 2013) and converted the resulting alignment to a codon alignment using pal2nal (Suyama et al., 2006). To reduce likely uninformative sites in the alignment we removed columns in which more than 70% of the sequences had gaps. The resulting codon alignment contained 2,292 nucleotide sites and was used to infer a phylogeny with IQ-TREE v.2.1.3 (Minh et al., 2020) under a GTR+F+I+R10 substitution model and node support was inferred with 10,000 Ultrafast bootstrap iterations (Hoang et al., 2018).

### 4.4 Recombination analysis

The 1,472 whole genome sequences (rotated to the start of ORF1 as described above) were aligned using mafft (–localpair option). These genomes contain extensive indel variation, resulting in many gappy regions in the alignment. Columns that consist primarily of gaps may misinform the recombination analysis, so all columns with nucleotides present for only one sequence (essentially representing insertions unique to a single genome) were removed from the alignment. The resulting alignment was examined for evidence of recombination using the Recombination Detection Program (RDP v.5.45) (Martin, Varsani, et al., 2021). Modules RDP (Martin and Rybicki, 2000), 3SEQ (Boni et al., 2007), GENECONV (Padidam et al., 1999), Chimaera (Posada and Crandall, 2001) and MaxChi (Smith, 1992) were performed followed by secondary scans with methods BootScan (Martin, Posada, et al., 2005) and SiScan (Gibbs et al., 2000). Sequences were assumed to be circular. Recombination events were accepted only when detected by at least by three of the methods used.

To assess potential clustering in presence or absence of detected breakpoints across the genomes’ length (hotspots or coldspots of recombination), we performed the breakpoint distribution test (BDT) and the recombination range test (RRT) also implemented in RDP5. The type sequence was set to the TTV reference genome (NC_002076), the window size was kept at 200 nucleotides and a total of 500 permutations were performed for each test.

### 4.5 Analysis of genomic features

The aforementioned ORF1 protein alignment (n=2,605) was used to explore the residue composition of the arginine-rich motif (ARM). The ARM region was defined as the N-terminal domain of each ORF1 until amino acid position 100 of the TTV reference genome (NC_002076) on the full ORF1 alignment. This region in the alignment matches the TTV ARM region as well as some upstream residues to capture arginine-rich regions extending further in other aligned sequences. Then, the number of arginines (R) and lysines (K) in that region were counted to represent the composition of positively charged amino acids in each virus’s putative ARM domain. Based on the distribution of R/K residue counts, it was observed that 265 ORF1s had less than 10 R/K residues in that part of the alignment (Figure S2). These sequences were assumed to be lacking their ARM domain and were excluded from downstream ARM composition analysis. The vast majority of remaining ORF1s had between 20 and 60 R/K residues in their ARM region (Figure S2). Three ORF1s had 82, 172 and 206 R/K residues respectively and were excluded from further analysis as outliers, likely due to sequence misalignment.

For the replication loop analysis, the reduced set of whole genome sequences (n=1,472) was used instead. Genome sequences classified into the three main clades (alpha, beta, gamma) based on the ORF1 phylogeny were aligned to their corresponding reference using mafft (–localpair) with the –keeplength option to remove insertions relative to the reference (Katoh and Standley, 2013) (alpha: TTV NC_002076 positions 3453-3460; beta: TTMV NC_014097 positions 2614-2621; gamma TTMDV NC_009225 positions 2946-2953). Members of the he and the novel X1-3 clades were aligned to the TTMV reference and manually inspected to confirm that the replication loop sequences were correctly aligned.

## Acknowledgements

We are grateful to Prof Fangqing Zhao for discussions about the sequencing of sample SRR2037085.

## Funding

SM was funded by an MRC Precision Medicine PhD studentship (MR/S502479/1). SL was funded by a MRC studentship. JH and RJO are funded by the MRC (MC_UU_1201412).

## Availability of data and materials

All raw data and associated code is available at https://github.com/spyros-lytras/anellome. All anellovirus contigs can also be accessed using BioProject PRJEB75196 [*upon publication*].

## Competing interests

The authors declare that they have no competing interests.

## Authors’ contributions

Conceptualisation: SM, SL. Data curation, Formal Analysis, Project administration, Investigation, Resources, Software, Validation, Visualisation and Writing - original draft SM, SL. Funding acquisition SM, SL. Methodology SM, SL. Writing - review & editing SM, JH, RJO, SL.

## Supplemental Material

### Supplementary figures

**Figure S1:**
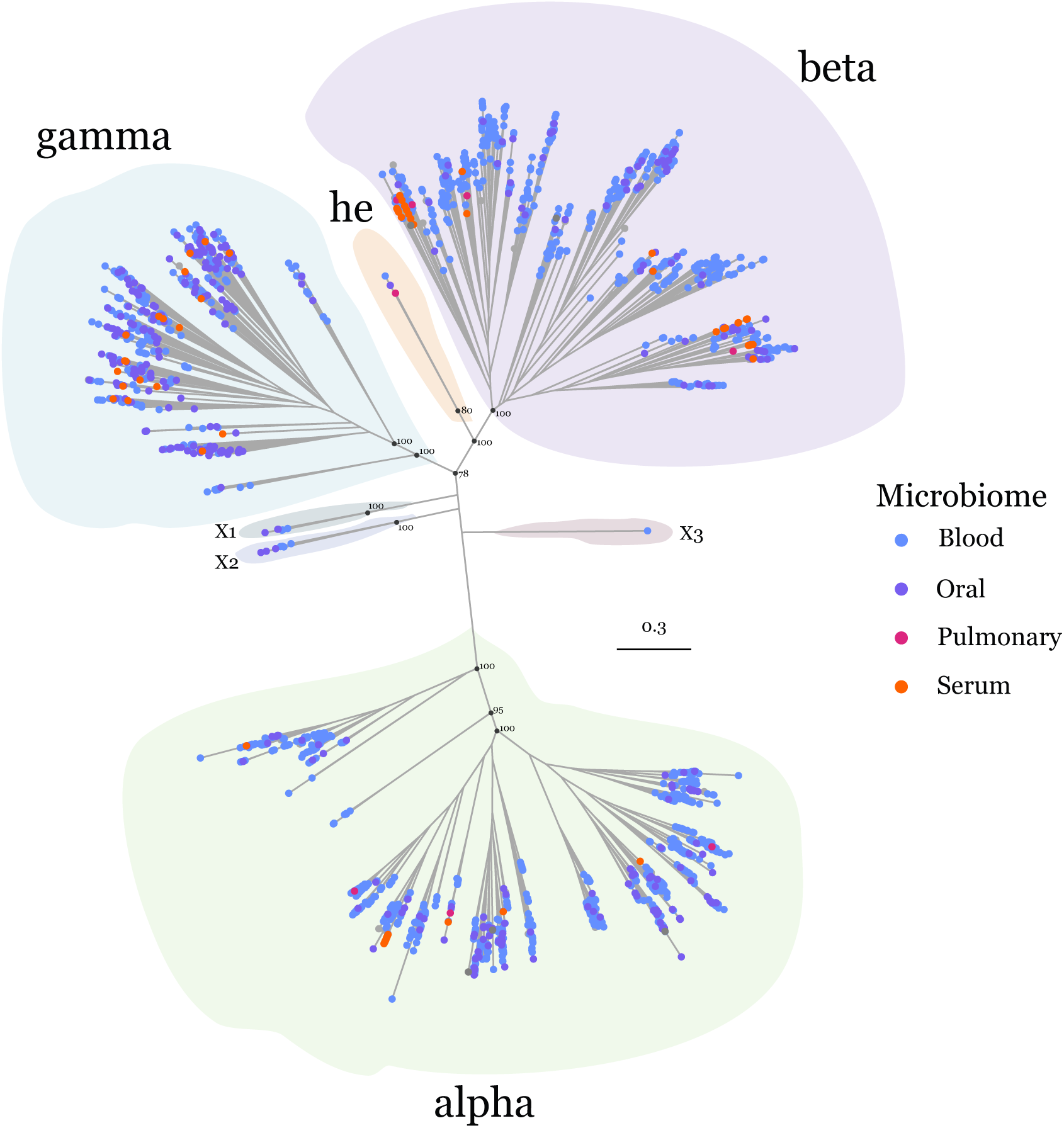
ORF1 phylogenetic tree annotated by the sampled microbiome. The original descriptions of the sample microbiomes were manually edited for consistency. Specifically, periodontal samples were put in the ‘Oral’ category; NT and pharyngeal swabs in the ’Pulmonary’ category; CSF and plasma samples in the ’Serum’ category. The tips of viruses associated with a microbiome category with less than 3 ORF1s (brain, stool, bone marrow, skin) and of viruses with no sample type information are coloured in grey and their categories are not displayed in the legend box. Ultrafast bootstrap support values are shown for key basal nodes. The clades of the phylogeny are shaded by their respective classification defined in Figure 1.

**Figure S2:**
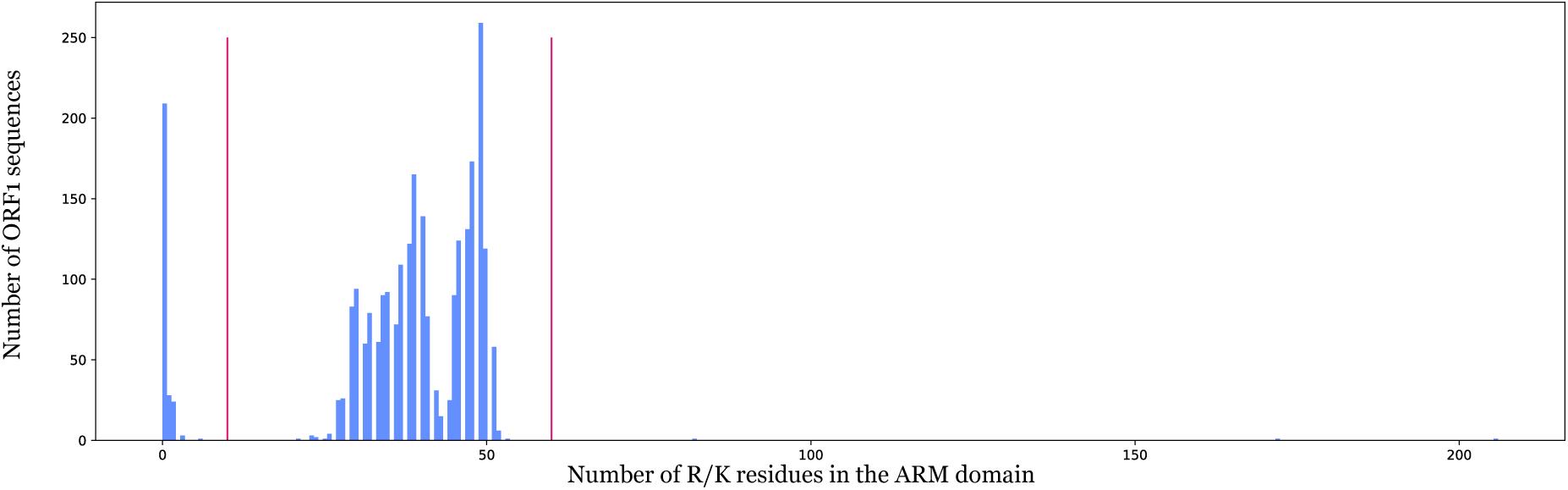
Distribution of the number of R and K residues in the ARM domain. Histogram representing the number of arginine (R) and lysine (K) residues in the determined ARM region of each analysed ORF1 protein sequence. The vertical pink lines indicate the determined cutoffs for the ARM domains used in downstream analysis (between 20 and 60 RK residues).

**Figure S3:**
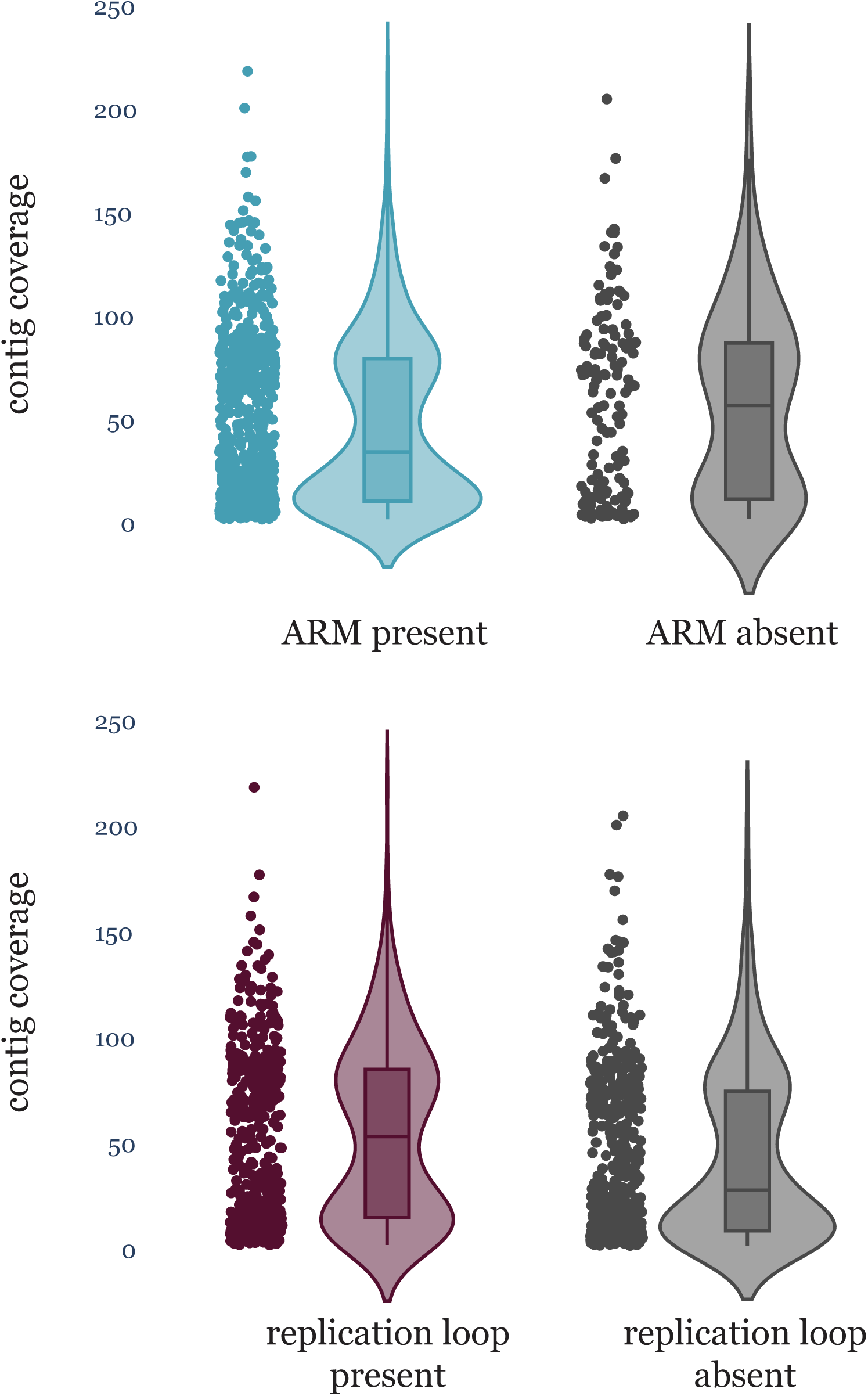
Comparison of contig coverage between genomes with and without key features. Distribution of the coverage of all anellovirus genome contigs assembled in this study separated by A) presence or absence of the ARM region (presence defined as having 20-60 RK residues, see Methods, Figure S2) and B) presence or absence of the replication loop region.

**Figure S4:**
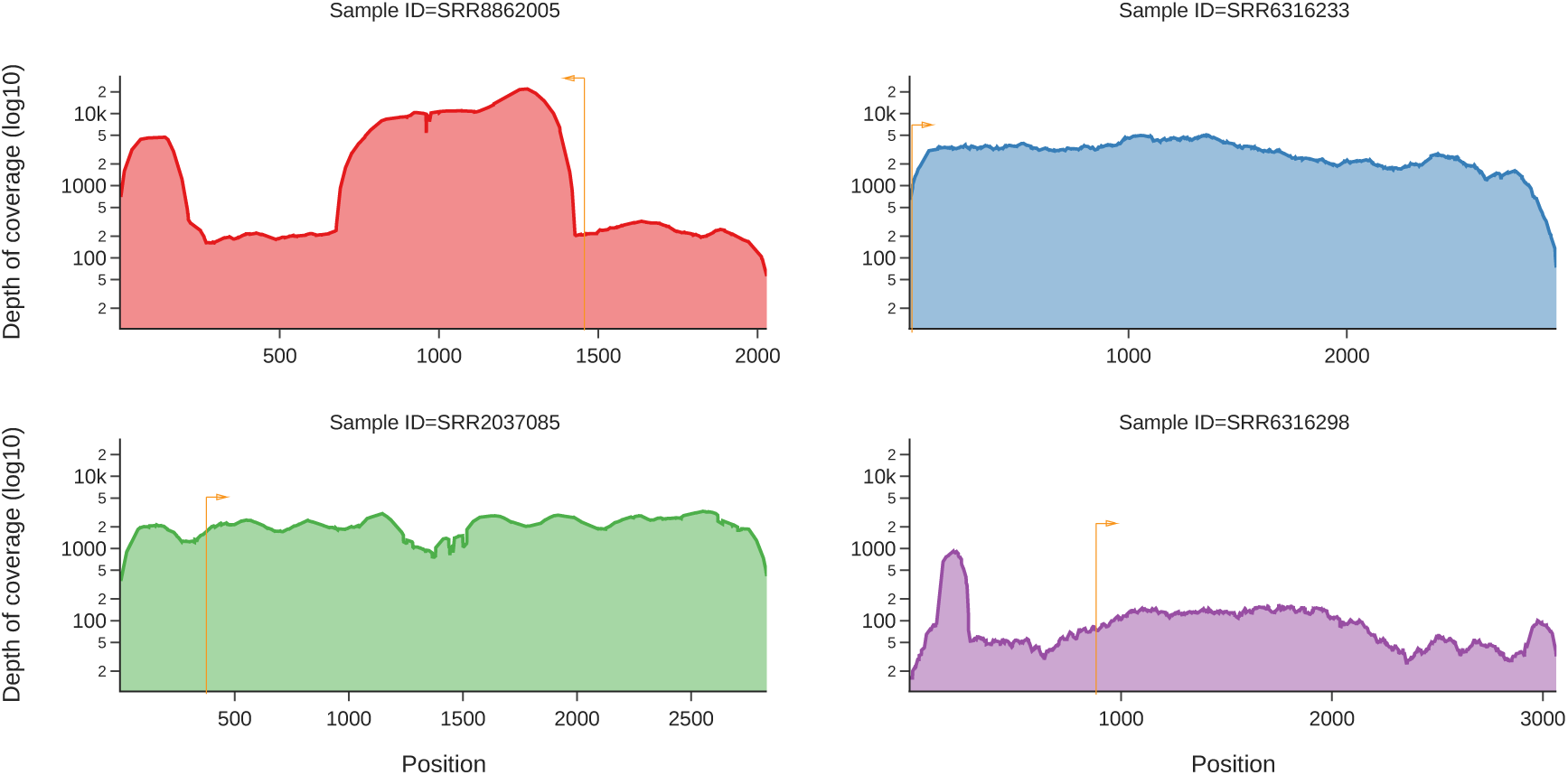
Per site coverage of high coverage contigs without ARM regions. Anellovirus sequences with high coverage (average depth >=100X) but missing ARM regions are analysed for depth of coverage (log value). To demonstrate that the absence of the ARM does not correlate with local dropouts in coverage, the coverage plots four different contigs from samples SRR8862005 (alpha, SRR8862005_NODE_16_length_2029_cov_205.534954, red, average depth: 4289.27), SRR6316233 (alpha, SRR6316233_NODE_15_length_2959_cov_142.645661, blue, average depth: 2873.52), SRR2037085 (alpha, SRR2037085_NODE_5287_length_2832_cov_109.884768, green, average depth: 2113.42), and SRR6316298 (alpha, SRR6316298_NODE_14_length_3064_cov_48.539050, violet, average depth: 110.764) are shown here. The coverage plots are calculated across assembled contigs that have not been rotated to the ORF1 start point, instead, the start point and coding direction of ORF1 is annotated by the orange line and arrow on each plot.

## Notes

https://github.com/spyros-lytras/anellome

### Competing Interest Statement

The authors have declared no competing interest.

